# Integrative analysis of DNA methylation suggests down-regulation of oncogenic pathways and reduced de-novo mutation in survival outliers of glioblastoma

**DOI:** 10.1101/302042

**Authors:** Taeyoung Hwang, Dimitrios Mathios, Kerrie L McDonald, Irene Daris, Sung-Hye Park, Peter C Burger, Sojin Kim, Yun-Sik Dho, Hruban Carolyn, Chetan Bettegowda, Joo Heon Shin, Michael Lim, Chul-Kee Park

## Abstract

The study of survival outliers of glioblastoma (GBM) can have important implications on gliomagenesis as well as in the identification of ways to alter clinical course on this almost uniformly lethal cancer type. However, current studied epigenetic and genetic signatures of the GBM outliers have failed to identify unifying criteria to characterize this unique group of patients. In this study, we profiled the global DNA methylation pattern of mainly IDH1 wild type survival outliers of glioblastoma and performed comprehensive enrichment analyses with genomic and epigenomic signatures. We found that the genome of long-term survivors in glioblastoma is differentially methylated relative to short-term survivor patients depending on CpG density: hypermethylation near CpG islands (CGIs) and hypomethylation far from CGIs. Interestingly, these two patterns are associated with distinct oncogenic aspects in gliomagenesis. The hypomethylation pattern at the region distant from CGI is associated with lower rates of de novo mutations while the hypermethylation at CGIs correlates with transcriptional downregulation of genes involved in cancer progression pathways. These results extend our understanding of DNA methylation of survival outliers in glioblastoma in a genome-wide level, and provide insight on the potential impact of DNA hypomethylation in cancer genome.

## Introduction

Despite advances in modern neurooncology, glioblastoma (GBM) continues to have a poor prognosis. Survival rates of adult GBM patients in the United States are quite low with 1-year, 2-year, 3-year, and 5-year relative survival rates estimated at 39.3%, 16.9%, 9.9%, and 5.5%, respectively (Ostrom et al. 2016). While the majority of GBM patients live no longer than 2 years, there is a subset of patients who live longer than 3 years and are classified as long-term survivors (LTS). This group of patients remains a puzzle to researchers in the field, as studies on clinical, radiological, histological, and molecular characteristics have yet to yield consensus regarding determinants of durable response to the current treatment (Scott et al. 1999; Burton et al. 2002a; Burton et al. 2002b; Shinojima et al. 2004; Krex et al. 2007; Martinez et al. 2007; Shinawi et al. 2013; Ma et al. 2015; Lu et al. 2016; Mock et al. 2016; Prasanna et al. 2016; Nakagawa et al. 2017; Peng et al. 2017). Classic genetic markers of favorable prognosis such as O-6-methylguanine-DNA methyltransferase (MGMT) promoter methylation or isocitrate dehydrogenase (IDH) mutation do not fully account for LTS-GBM (Hartmann et al. 2013; Gerber et al. 2014; Amelot et al. 2015; Millward et al. 2016; Sarmiento et al. 2016; Smrdel et al. 2016). Review of the literature reveals a report of concurrent gain of chromosomes 19 and 20 as a favorable prognostic factor for a subset of LTS-GBM that did not show IDH mutation-related DNA methylation pattern called ‘Glioma CpG Island Hypermethylator Phenotype (G-CIMP)’ (Geisenberger et al. 2015). However, multiple studies revealed no distinctive DNA copy number changes in LTS-GBM (Hartmann et al. 2013; Gerber et al. 2014; Reifenberger et al. 2014). Also, efforts to identify specific gene expression profiling patterns for LTS-GBM failed to uncover consistent features (Gerber et al. 2014; Reifenberger et al. 2014; Geisenberger et al. 2015). These results suggest that there is little chance to discover a single genetic or epigenetic biomarker that can define the LTS-GBM group, emphasizing the importance of comprehensive understanding of molecular signatures in LTS-GBM. In fact, a recent integrated genomic analysis comparing LTS and short-term survivors(STS) GBM showed that multiple genetic and epigenetic mechanisms are involved in divergent molecular features between the two extremes of the survival spectrum (Peng et al. 2017).

Although there have been numerous genome-wide studies for DNA methylation in brain cancer, most of them have largely focused on promoter regions and CpG islands (CGIs) in identifying aberrant methylation patterns or in classifying GBM patients due to its readiness of interpretation in terms of transcriptional regulation (Brennan et al. 2013; Mack et al. 2016). However, DNA methylation outside promotor-associated CGIs also presents distinctive signatures in tumors and has significant effects on regulation of genes in oncogenic pathways. For example, DNA methylation of the CpG sites in gene body is known to be a major cause of cytosine to thymine transition mutations, which can cause oncogenesis (Jones 2012). In addition, DNA methylation in the gene body is known to stimulate transcription elongation unlike the methylation of CGIs in promoter region, which is associated with gene expression silence (Jones 2012). Moreover, there is a genome-wide crosstalk, not limited to genic region, between DNA methylation and histone modifications (Malzkorn et al. 2011; Rose and Klose 2014). One good example is that trimethylation of histone H3 lysine 9 (H3K9me3) is required for DNMT3B dependent de novo DNA methylation (Rose and Klose 2014). Therefore, unbiased and integrative analysis of DNA methylation pattern across the whole genome is necessary to define GBM of exceptional clinical course.

In the present study, we compared genome-wide DNA methylation profiles of IDH wild-type (IDH WT) GBM patients who lived longer than 3 years (n=17, LTS-GBM) with patients who lived less than 1 year (n=12, STS-GBM) and performed integrative analysis of differential DNA methylation signatures across the whole genome to understand their functional implications on epigenetic aspects of GBM. We found that there are striking differences in methylation profiling signatures between LTS- and STS-GBMs in two independent cohorts. The most significant difference was found in CGIs and the region distant from CGI, called “open sea” where LTS-GBM showed hypermethylation and hypomethylation, respectively. The hypermethylation in LTS-GBM is preferentially found in promoter region of high CGI density and is enriched with histone marks of active transcription such as the acetylation of histone H3 lysine 27 (H3K27ac). The hypermethylation is associated with downregulation of transcription and the genes associated with hypermethylation in LTS-GBM are involved in oncogenic pathways including cellular proliferation and cell-to-cell attachment. On the contrary, H3K9me3 is the chromatin mark that was enriched in the hypomethylated sites of open sea in LTS-GBM. We found that the occurrence of somatic mutations in the sites around the open sea correlate with higher DNA methylation, implying that hypomethylation in LTS-GBM is associated with lower regional somatic mutation rates.

## Results

### Differential methylation profile between LTS- and STS-GBM

The statistical tests comparing the methylation status of 17 IDH WT LTS-GBM with those of 12 IDH WT STS-GBM identified 162,136 autosomal sites showing significant differences (see methods, FDR<0.01) in methylation levels (Fig. 1a). They generally show moderate differences in methylation level (mean differences of beta values<0.3, supplementary fig. 1). Differentially methylated sites are evenly distributed throughout the chromosomes (Supplementary fig. 2). Interestingly, the significant sites tend to have higher methylation level in LTS when they are closer to CGI or transcription start site (TSS), while lower methylation was observed in the distant region of open sea from island or TSS in LTS compared with STS (Figs. 1b and 1c).

**Figure 1.**
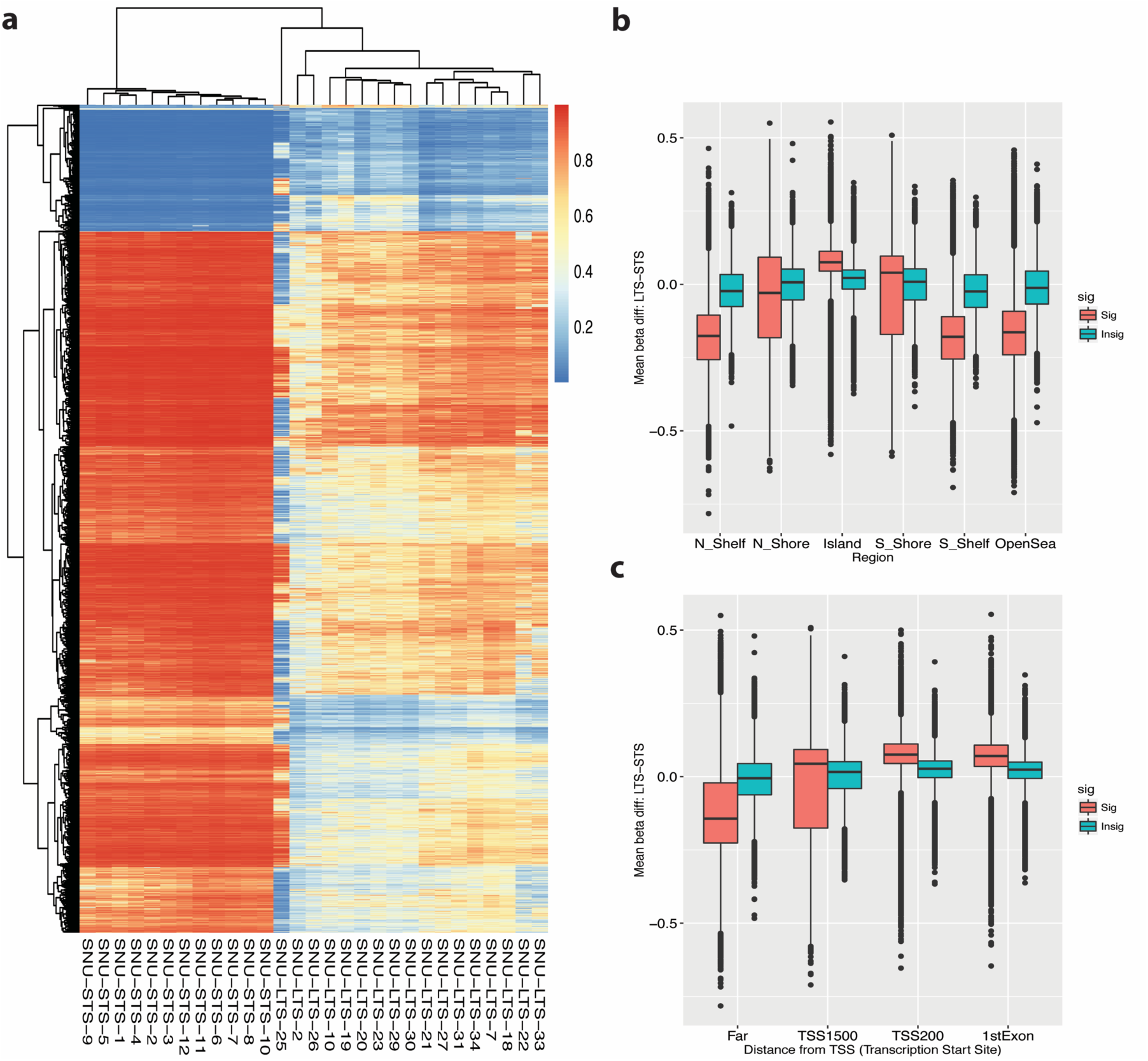
Differentially methylated sites between LTS-GBM and STS-GBM. **(a)** Heatmap of DNA methylation levels measured as beta values: color gradients from blue to red correspond to beta values from 0 to 1. Hierarchical clustering was performed for both of glioblastoma patients (columns) and the differentially-methylated sites between LTS-GBM and STS-GBM (rows). The most significant sites were used for this plot for convenience (−log_10_(FDR)< 5: 11823 sites). **(b)** Distribution of mean differences of beta values between LTS-GBM and STS-GBM across the sites categorized by their locations from the closest CGI (see methods). The group of selected significant sites are denoted by red while the other sites are described by green. **(c)** Same as (b) except that categorization of sites were done by distances from the nearest transcription start site (see methods).

**Supplementary figure 1.**
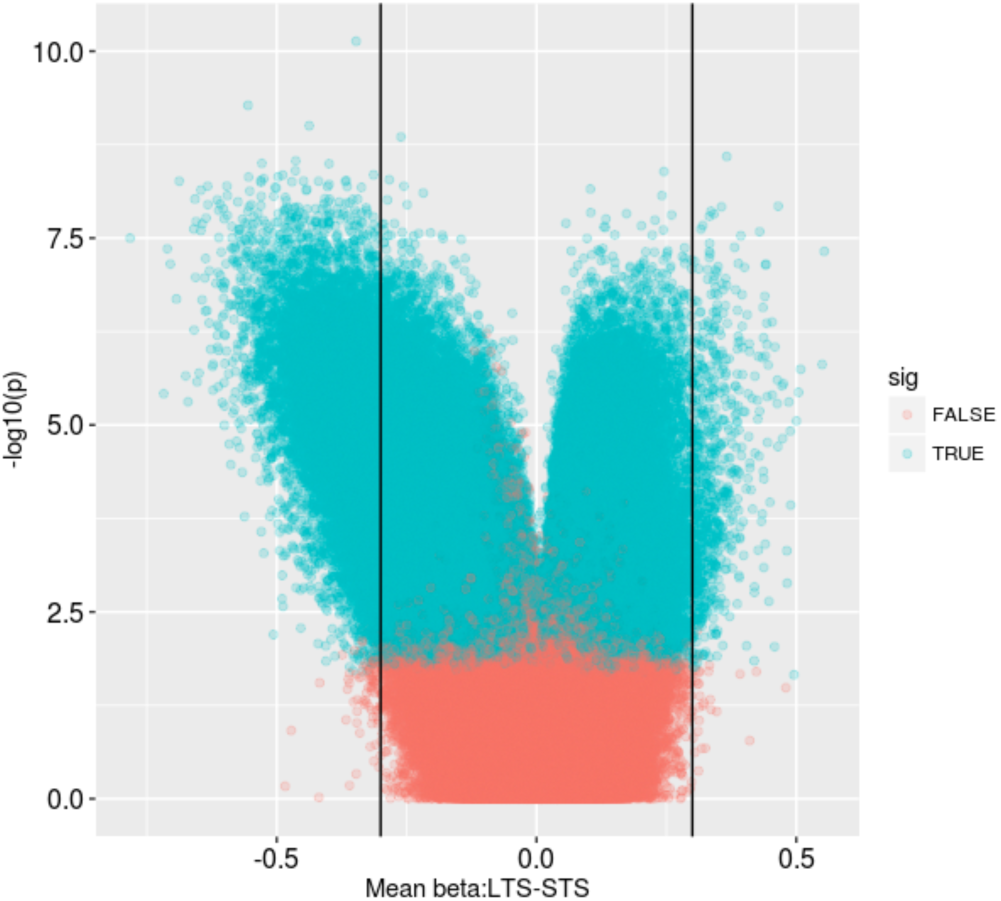
Volcano plot of differences between STS-GBM and LTS-GBM in DNA methylation level (beta). Each dot is a single CpG site measured by a probe. X axis is mean difference of beta values between STS-GBM and LTS-GBM groups. Y axis is statistical significance. Red dot indicates significant sites while green dot corresponds to a site without statistical significance. Two vertical lines denote mean differences of −0.3 and 0.3 respectively.

**Supplementary figure 2.**
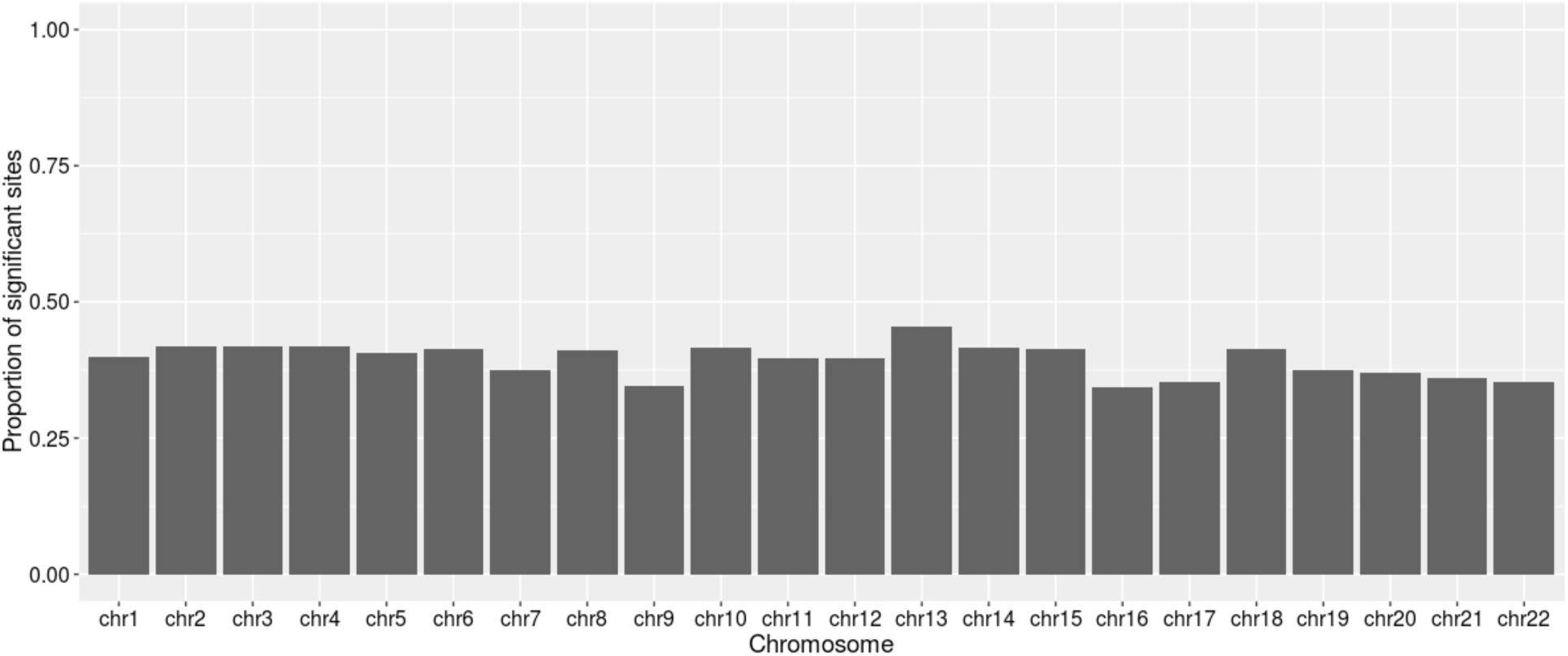
Proportion of significant sites in comparison of DNA methylation levels (beta) between STS-GBM and LTS-GBM. X axis is autosomal chromosome and Y axis is proportion of significant sites in a given chromosome.

In order to delineate the differential pattern in greater detail, we investigated the association of the identified patterns with histone modifications by using the representative sites of the two groups of significant sites: the sites in CGI that are hypermethylated and the sites in open sea that are hypomethylated. Specifically, the two groups of genomic sites were constructed by selecting the sites whose mean difference of beta values between LTS-GBM and STS-GBM was greater than 0.2 in each region, and were correlated with the genomic regions of histone marks in the ENCODE consortium dataset (see methods). The two groups of the significantly different methylation sites showed distinct enrichment of histone marks compared to the insignificant methylation sites (Fig. 2). For example, H3K27ac was most enriched with hypermethylated sites in island region, however, this was underrepresented in the hypomethylated sites in the open sea region. On the contrary, H3K9me3 was mostly enriched in the significant sites in the open sea region while it was depleted in the island region. These results imply that our comparison between LTS-GBM and STS-GBM identifies differential DNA methylation signatures with regulatory potential. In fact, both histone marks of H3K27ac and H3K9me3 are known to be related with DNA methylation (Charlet et al. 2016).

**Figure 2.**
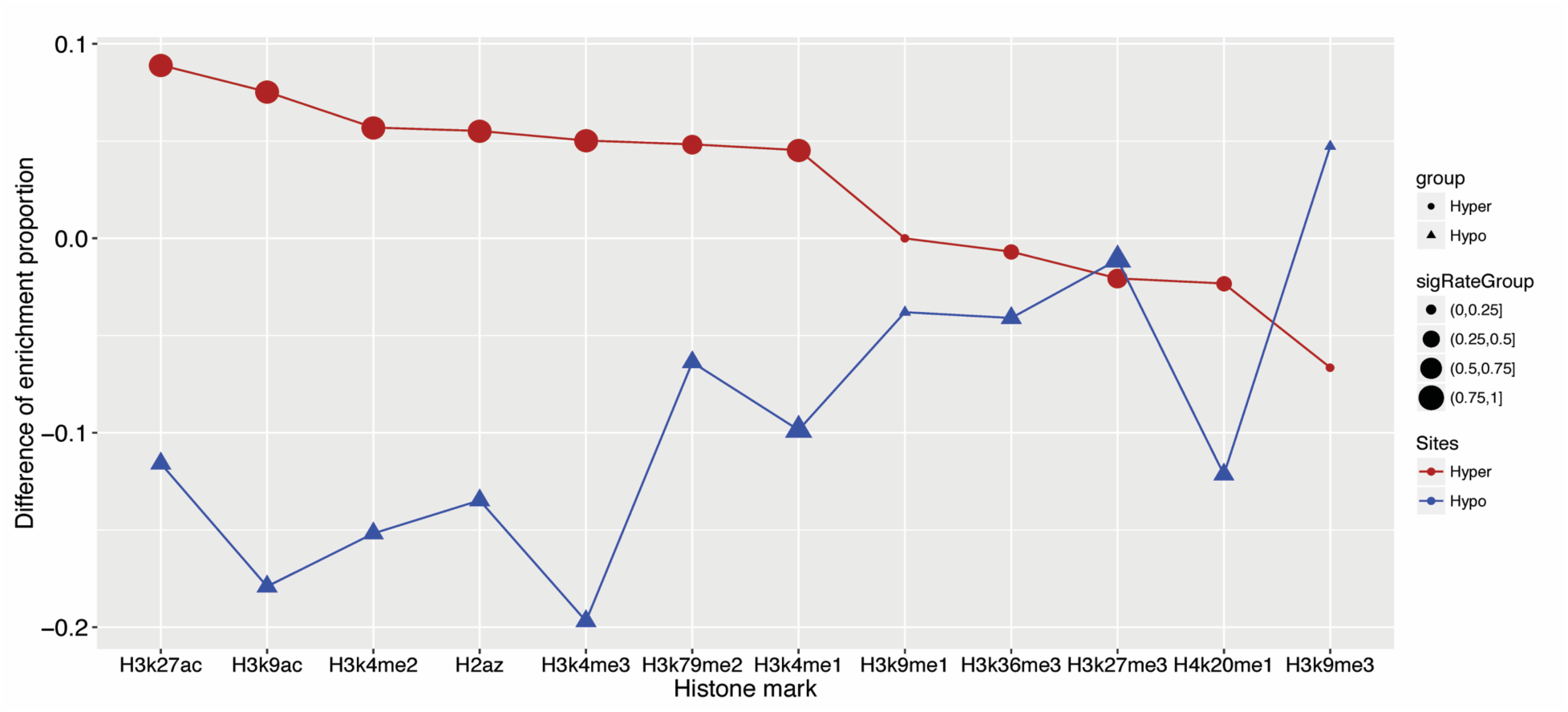
Enrichment of differentially-methylated sites in regulatory histone marks. Enrichment pattern of differentially-methylated sites in histone marks. For a given histone mark (x axis), the significant differential sites between STS-GBM and LTS-GBM are tested for their enrichment. Y axis describes the differences of enrichment proportion between the significant differential sites and the insignificant sites. The red denotes hypermethylated sites in island while the blue describes hypomethylated sites in open sea. The size of dot indicates the range of enrichment proportion of the significant differential sites. We grouped the proportion to 4 regions for clear visualization: 0∼0.25, 0.25∼0.5, 0.5∼0.75, 0.75∼1.

### LTS-GBM-specific hypermethylation in CGI (“island”)

Enrichment of histone marks of active transcription such as H3K27ac and H3K9ac in hypermethylated sites in island suggests their potential in transcriptional regulation. These sites are also enriched in promoter regions (Supplementary fig. 3a). In fact, it is well known that promoter region is represented by histone marks of H3K9ac and H3K27ac. As the role of DNA methylation in the promoter region is known to repress transcription, we tested whether DNA methylation of these sites is associated with varying gene expression levels. Specifically, the distribution of correlations between DNA methylation and gene expression across GBM samples in the TCGA consortium were compared between these sites and the random genic sites in the CGI region. The enrichment of negative correlation between methylation and expression was observed for the sites showing the hypermethylated sites relative to the non-significant sites in island (Fig. 3a). Furthermore, we confirmed that the DNA methylation level (retrieved from TCGA) tends to negatively correlate with H3K27ac (obtained from Lin et al. 2012) in U87MG cell line at these hypermethylated sites (Fig. 3b, coefficient of linear regression: −0.16 with N=1298, p-value<10^−15^). We performed gene ontology (GO) analyses to understand genes regulated by hypermethylation in LTS-GBM. Genes associated with hypermethylated sites in islands are enriched with the gene ontology terms related to cancer progression such as cellular proliferation and cellular attachment (Fig. 3c and Supplementary table 2). These results imply that hypermethylation in LTS suppress gene expression in tumor progression pathways possibly mediated through histone marks of active transcription around the CGIs.

**Figure 3.**
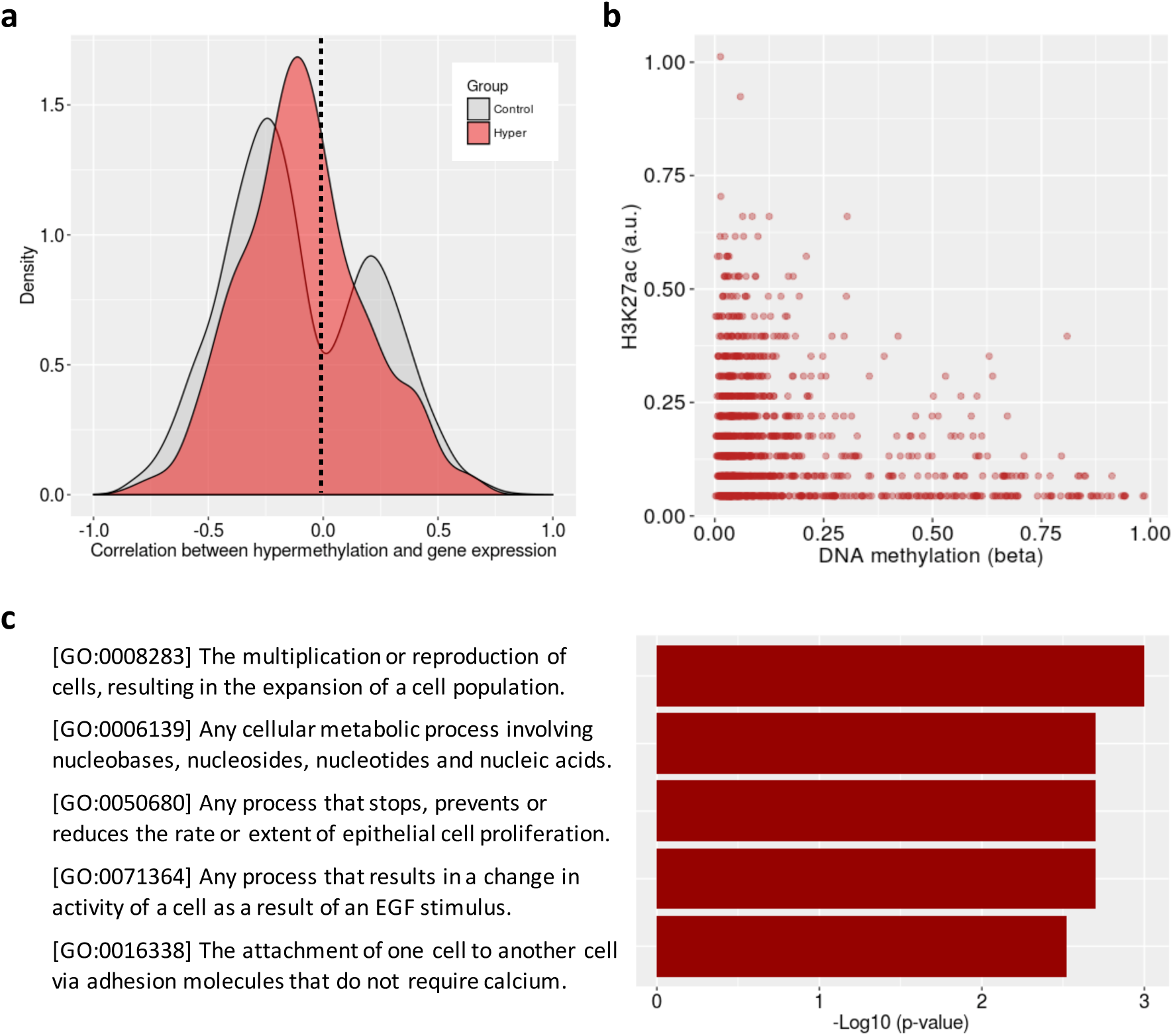
Transcriptional regulatory potential of hyper-methylated sites. **(a)** Distribution of Pearson correlation coefficients between DNA methylation level (beta) and gene expression measured by RNA-seq (FPKM) across GBM samples in TCGA. The red describes the selected hyper-methylated sites in this study while the gray shows the other sites associated with the island, **(b)** The relation between the ChlP-seq signal of H3K27ac and the DNA methylation level (beta) for the hyper-methylated sites in U87MG cell line. Each dot denotes the site, **(c)** The top five gene ontology terms (category: biological process) highly enriched with the genes associated with the selected hyper-methylated sites.

**Supplementary figure 3.**
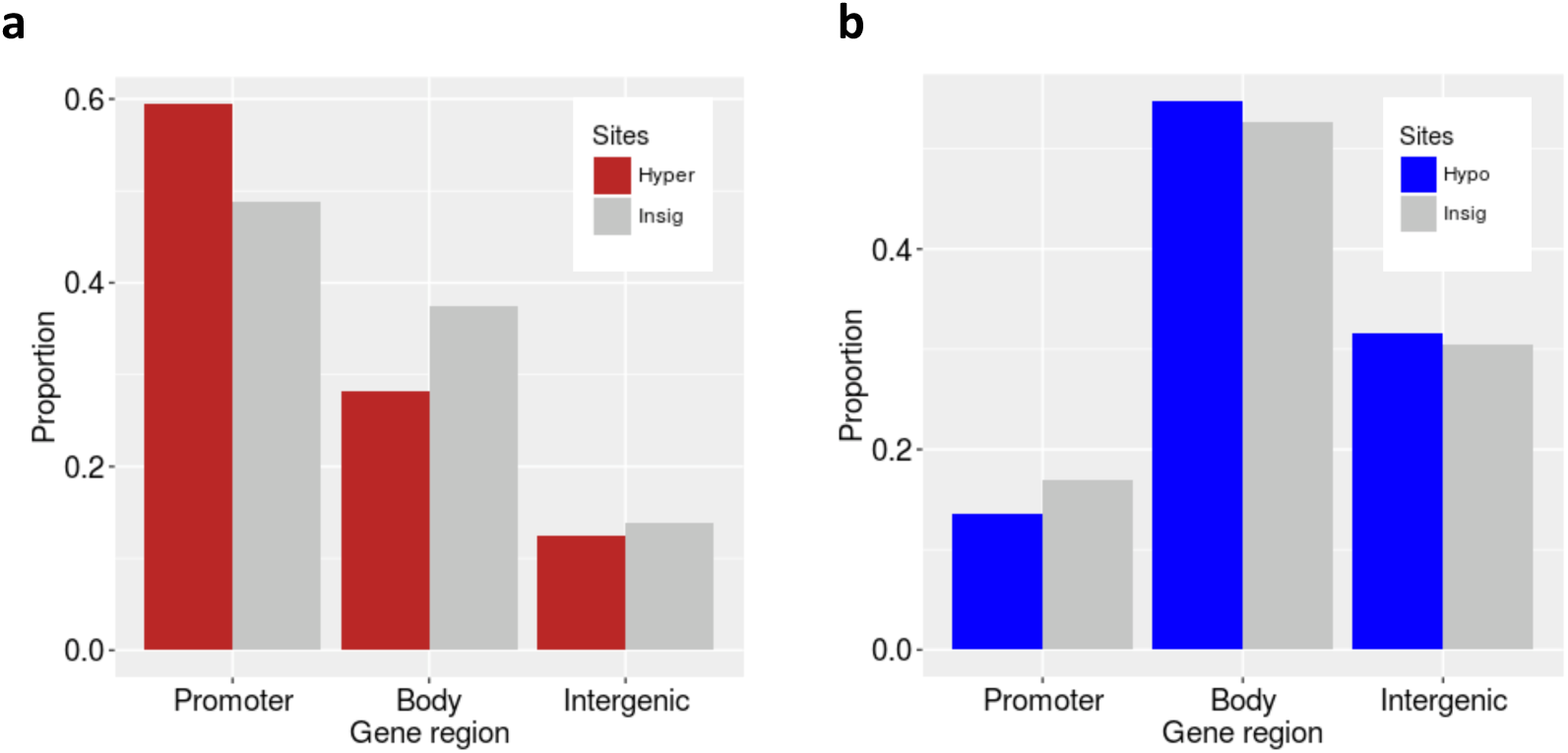
Enrichment of significant differential sites in gene regions. **(a)** Hypermethylated sites in island **(b)** Hypomethylated sites in open sea. Promoter is enriched with the hypermethylated site relative to gene body while gene body is enriched with the hypomethylated site relative to promoter (p-values<10^−15^ from Fisher exact tests in both cases).

### LTS-GBM-specific hypomethylation in open sea (distant region from CGI)

Although the hypomethylated sites in the open sea were enriched with a histone mark of heterochromatin, H3K9me3, their enrichment with specific gene region is not as obvious as the hypermethylated sites (Supplementary fig. 3b). We only found a modest enrichment in gene body region for the hypomethylated sites. Also, the hypomethylated sites showed the bimodal distribution of positive and negative correlation coefficients with the associated mRNA expression, weakening its consistent global regulatory potential for transcription (Supplementary fig. 4). Furthermore, the hypomethylation-associated genes did not identify GO terms related to cancer (Supplementary table 3). We hypothesized that hypomethylation in LTS-GBM has less implication on gene activity of oncogenic pathway.

Previous studies suggest that epigenetic modifications of the genome can affect the mutational rate in the human genome (Makova and Hardison 2015). We hypothesized that hypomethylation in the open sea might affect local mutational rate and tested our hypothesis for exon regions in a genome-wide way using the available whole exome sequencing data for TCGA GBM samples. Specifically, we compared DNA methylation levels near somatic mutation sites between the hypomethylated sites and the insignificant sites in open sea. It turned out that if somatic mutations were observed around the identified hypomethylated sites in LTS-GBM, the sites tend to be more methylated in TCGA GBM samples. However, the sites not specific to LTS-GBM in open sea do not show such a bias of DNA methylation (Fig. 4a). We also confirmed that DNA methylation is positively correlated with H3K9me3 at our hypomethylated sites in H1 cell line using the ENCODE dataset that provides both measurements of H3K9me3 ChIP-seq and 450k bead array (Fig. 4b, coefficient of linear regression: 0.29 with N=24372, p-value<10^−15^). These results imply that hypomethylation featured in open sea in LTS-GBM may contribute to lower mutational rates, which involved with changes in H3K9me3, a histone mark of heterochromatin context.

**Figure 4.**
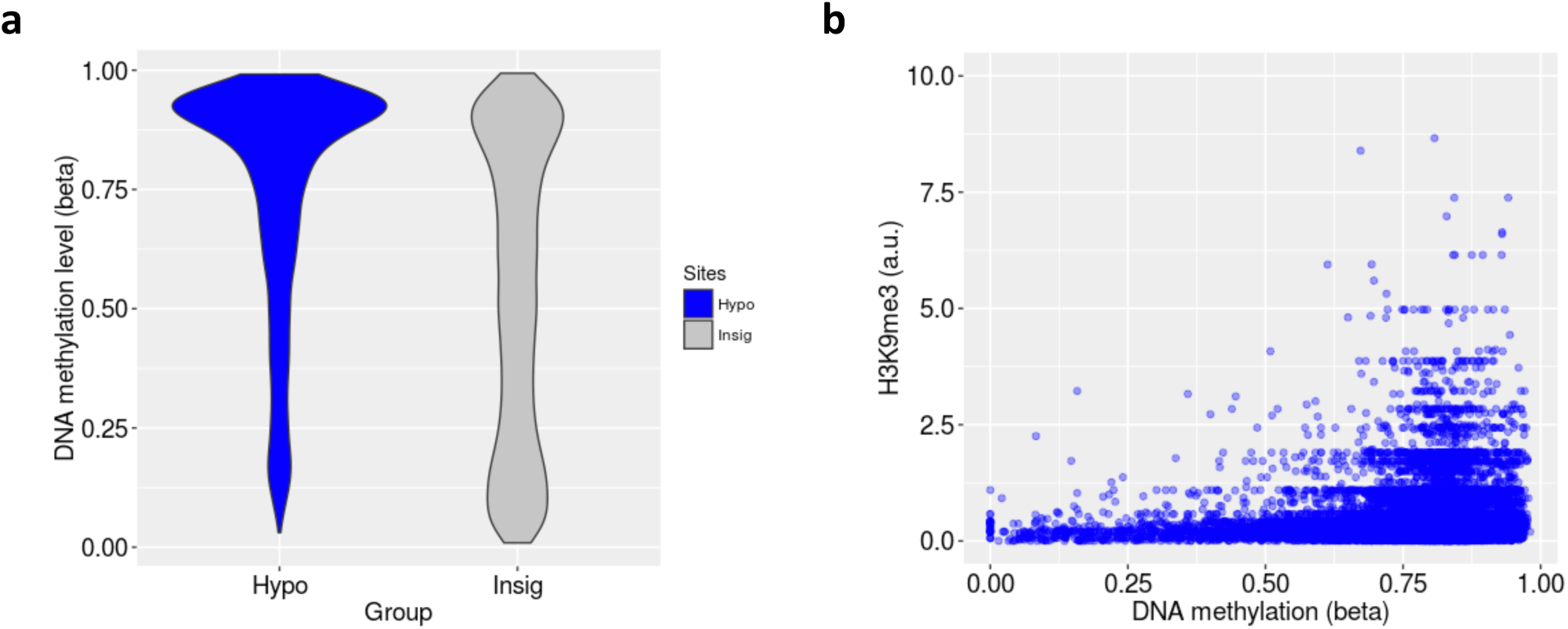
Bias of DNA methylation level near local somatic mutations in open sea (far-CGI region). **(a)**DNA methylation level around somatic mutations found in 135 TCGA GBM samples by whole exome sequencing. The hypo-methylated sites in LTS-GBM (N=5875) were compared with the insignificant sites in open sea (N=21630). **(b)** The relation between the ChIP-seq signal of H3K9me3 and the DNA methylation beta for the hyper-methylated sites in H1 cell line. Each dot denotes the site.

### Validation of differential methylation in an independent LTS-GBM cohort

Finally, we asked if our identification of hypermethylation in CGIs and hypomethylation in open sea is reproducible in independent cohorts of LTS-GBM. Hierarchical clustering was performed for two independent test sets of IDH1 WT GBM samples (TCGA and ‘Australian’, see methods with supplementary table 1) with DNA methylation level at the identified hyper or hypomethylated sites (Fig. 5a). First, all of 9 Australian LTS samples show similar pattern of hyper and hypomethylation. Second, 38 out of 39 STS in the TCGA database replicated our pattern while only 3 LTS-GBM samples were not accounted for by our pattern.

We also attempted to assign scores to each test sample in terms of hypermethylation and hypomethylation in order to predict LTS-GBM. The simple arithmetic average of beta values in each pattern is assigned to a sample. We predicted a sample as LTS-GBM when its mean beta values for both of hypermethylation and hypomethylation are greater than 0.2, otherwise it was called as STS-GBM. As a result, 9 out of 12 LTS-GBM were predicted correctly (sensitivity: 75%) and 38 out of 39 STS samples are correctly recalled (specificity: 97%) (Fig. 5b).

**Figure 5.**
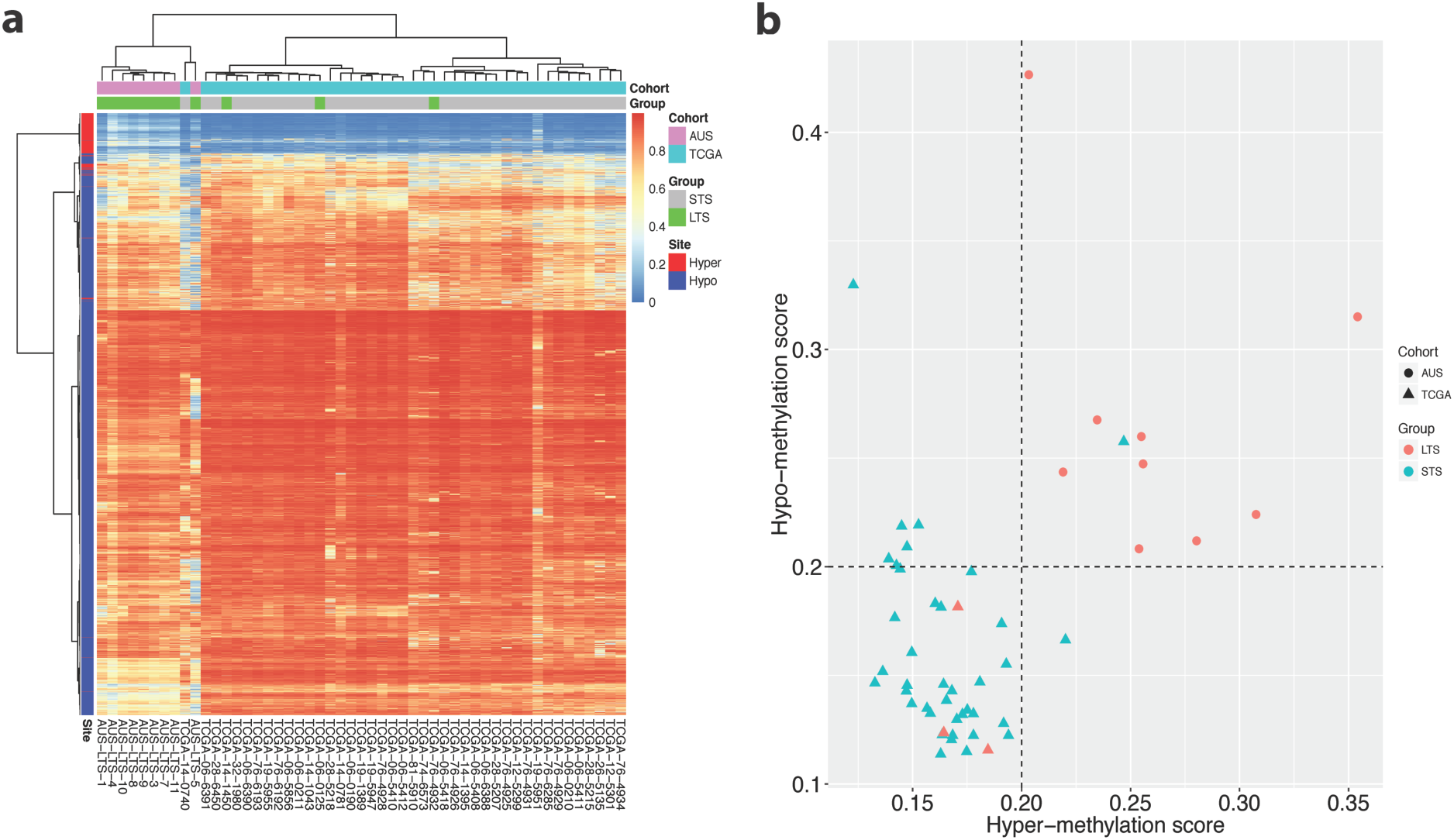
Hyper- and hypo-methylation in two independent cohorts (TCGA and ‘Australian’). **(a)**Heatmap of DNA methylation levels measured as beta values: hierarchical clustering was performed for both of samples (columns) and the sites (rows) either hyper- or hypo-methylation identified in the discovery cohort (“SNU”). The color gradients from blue to red correspond to beta values from 0 to 1. **(b)** Summary scores in terms of hyper- and hypo-methylation for each sample in two test cohorts: Each sample, denoted by a dot is assigned to two simple arithmetic averages (x and y axes values) of beta values in hyper- and hypo-methylation sites. The dashed line indicates 0.2 as a decision threshold for LTS-GBM.

## Discussion

Epigenetic aberrations are increasingly regarded as a gateway to neoplastic transformation in gliomas (Mack et al. 2016). In particular, a recent study showed that DNA methylation was the strongest predictor of prolonged survival in GBM compared to any of clinical variables, RNA expressions for mRNA/miRNA, and the available genomic data including germline/somatic point mutation and copy number variation (Lu et al. 2016). They found the importance of DNA methylation based on the statistical analyses for clinical data and multimodal molecular profiles of 44 patients (7.4%) who lived longer than 3 years among 591 GBM patients from TCGA dataset. However, the effect of DNA methylation is often confounded with genetic perturbation. For example, although G-CIMP signatures were found to be a favorable prognositic marker of GBM, the majority of them overlap with IDH mutation (Parsons et al. 2008; Turcan et al. 2012). However, IDH mutation cannot explain all the cases of LTS-GBM and there is a subset of LTS-GBM patients with wild-type IDH (Gerber et al. 2014; Reifenberger et al. 2014; Amelot et al. 2015; Sarmiento et al. 2016). Therefore, it is important to evaluate the effect of DNA methylation in LTS-GBM after controlling genetic background such as IDH1 mutation. There have been several studies identifying DNA methylation signatures specific for IDH1 WT LTS-GBMs. Mock et al. compared global DNA methylation profiling using Methyl-CpG-Immunoprecipitation in 14 LTS and 15 STS-GBM patient samples with IDH1 wild-type, and found that hypermethylation of multiple CpGs mapping to the promoter region of *LOC283731* correlated with improved patient outcome (Mock et al. 2016). Zhang et al. analyzed methylation profiles of 13 LTS and 20 STS-GBM patients using Illumina Infinium Human Methylation 27K Bead-Chips (Zhang et al. 2013). They identified the promoter methylation in *ALDH1A3* is a prognostic biomarker in a IDH1 wild-type and unmethylated MGMT promoter GBM sample. However, these studies only focused on DNA methylation in promoter regions and did not provide comprehensive understanding of landscape picture of DNA methylation signatures in LTS-GBM.

In the present study, we pursued the genome-wide understanding of DNA methylation patterns specific to IDH1 WT LTS-GBM. We showed that LTS-GBM, compared with STS-GBM, is characterized by hypermethylation in the CGI regions and hypomethylation at open sea throughout the genome region which were replicated in the two independent cohorts of LTS-GBM. It is well known that methylation of CGIs at promoter region is linked to silence of gene expression, and our results were consistent with this idea and previous findings of hypermethylation in LTS-GBM. However, hypomethylation of open sea region in LTS-GBM is poorly appreciated so far. It is not compelling to understand this pattern in terms of gene activity since our GO analysis showed that the genes are not generally related with cancer. Also, as genome-wide hypomethylation in cancer is known to be ubiquitous feature of carcinogenesis (Ehrlich 2009), the hypomethylation pattern of LTS-GBM relative to STS-GBM seems to be enigmatic. However, a previous study showed that methylated cytosine bases are prone to mutation by spontaneous deamination to thymine (Rideout et al. 1990). Additionally, there is evidence linking levels of regional mutation density of cancer cells with the heterochromatin-associated H3K9me3 that is a sole histone mark enriched with our hypomethylated sites (Schuster-Bockler and Lehner 2012). Our analyses of exome-sequencing of GBM showed that local mutations tend to happen at the identified sites in open sea when they have higher DNA methylation levels. This suggests that significantly lower methylation around the identified sites in LTS-GBM can reduce the risk of de novo mutation contributable to oncogenesis possibly in the absence of proper DNA repair mechanisms in GBM.

Altogether, our results provide comprehensive understanding of DNA methylation in survival outliers of glioblastoma (Fig. 6). But elucidating the mechanism of how hypomethylation in LTS-GBM is contributing to better survival relative to STS-GBM remains to be determined by further studies. Our results call more attention to dual aspects of DNA hypomethylation in open sea that have implications on both oncogenic contribution and survival benefits of patients.

**Figure 6.**
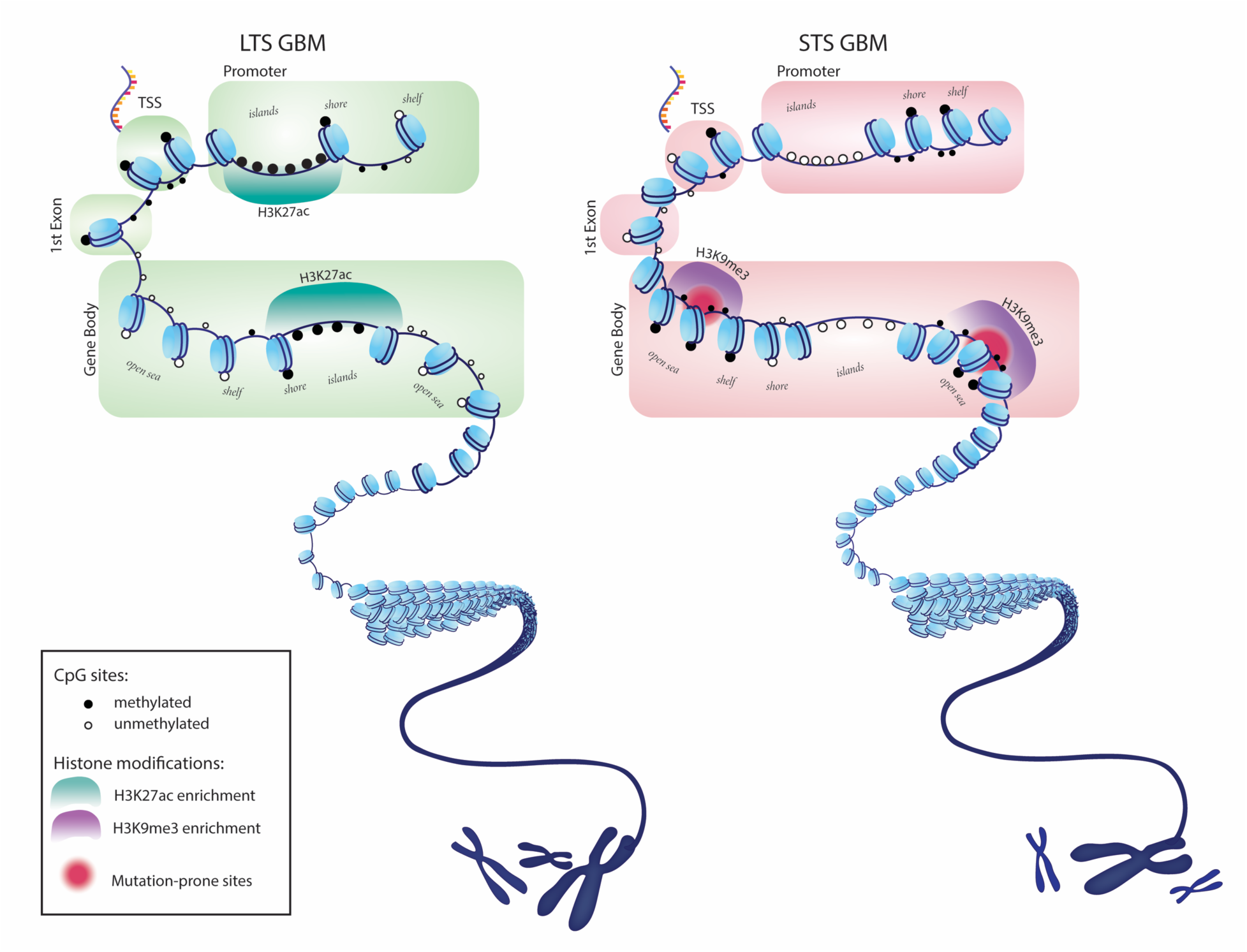
Genome-wide DNA methylation pattern of glioblastoma. The genomes of long-term survivors in glioblastoma are differentially methylated relative to short-term survival patients depending on CpG density: hypermethylation near CpG islands(CGIs) and hypomethylation far from CGIs (open sea). The hypermethylation at CGIs frequently occurs around regions with histone marks of active transcription such as H3K27ac, correlating with downregulation of gene expression in cancer progression pathways. The hypomethylated region at open sea are enriched with a histone mark of heterochromatin, H3K9me3. The rate of de novo mutation is high in this region when it is methylated, implying survival advantage of hypomethylation of the region in glioblastoma. In the figure, we highlighted genic regions such as first exon and gene body to emphasize potential effect of perturbed DNA methylation in glioblastoma.

## Methods

### Patient samples

Survival data of patients with histologically confirmed GBM at Seoul National University Hospital, Korea between 2000 and 2010 were obtained by retrospective review of patients’ charts, and from National Cancer Registry survival database of Korea. We identified 34 out of 429 newly diagnosed patients who lived longer than 3 years, and 17 patients were finally selected for discovery cohort of LTS-GBM group after central histological review. Central histological review included a complete agreement of histological diagnosis between two neuropathologists (S.H.P. and P.B.) on their independent slide review, and no evidence of IDH1 mutation on immunohistochemical evaluation. For the comparison, 12 GBM patients who lived less than 1 year in spite of standard care were chosen for the STS-GBM group. Patient clinical characteristics are summarized in the supplementary table 1.

For the validation cohort, 10 GBM samples were obtained from patients treated at Prince of Wales Clinical School, Australia between 2004 and 2009 who lived longer than 3 years (Supplementary table 1). This cohort contained 5 wildtype IDH1, 1 mutated IDH1, and 4 patients with unknown IDH1 status. We also collected IDH1 WT TCGA samples that provide raw data of Illumina’s Infinium Human Methylation450K BeadChips: 3 LTS and 39 STS samples.

This study was approved by the institutional ethics committees.

### DNA extraction

DNA was extracted by standard methods for formalin-fixed paraffin-embedded tumor tissue. The QIAamp DNA FFPE Tissue Kit (Qiagen) was used to isolate and purify DNA from formalin fixed, paraffin embedded tissue. The protocol suggested by the manufacturer was used to isolate DNA. The quality and quantity of DNA was assessed both by Nanodrop and Bioanalyzer technology. The isolation of DNA from all clinical samples was performed by the Genetics Core Store of Johns Hopkins School of Medicine.

### Genome-wide DNA methylation mesurement

Bisulfite-converted DNA was analyzed using Illumina’s Infinium Human Methylation450K BeadChips according to the manufacturer’s manual. DNA bisulfite conversion was carried out using EZ DNA Methylation Kit (Zymo Research) by following manufacturer’s manual with modifications for Illumina Infinium Methylation Assay. Briefly, 400 ng of genomic DNA was first mixed with 5 ul of M-Dilution Buffer and incubate at 37°C for 15 minutes and then mixed with 100 ul of CT Conversion Reagent prepared as instructed in the kit’s manual. Mixtures were incubated in a thermocycler with 16 thermal cycles at 95°C for 30 seconds and 50C for one hour. Bisulfite-converted DNA samples were loaded onto 96-column plates provided in the kit for desulphonation and purification. Concentration of eluted DNA was measured using Nanodrop-1000 spectrometer.

### Preprocessiong of DNA methylation array data: filtering probes and intra-sample normalization

We filtered the following probes in the Illumina Infinium HumanMethylation450 BeadChip: i) probes with a detection p-value is above 0.01 in more than 5% of samples. According to Illumina, the detection-p-value of a probe is estimated by comparing the measured CpG site intensity (the sum of methylated probe intensity and unmethylated probe intensity) with the intensities of negative control probes., ii) the probes whose bead counts are less than 3 in more than 5% of samples, iii) probes with SNP sites, iv) probes whose sequences align to multiple locations in the reference genome, v) probes associated with the sex chromosomes. The filtering was done by using R package ChAMP (Morris et al. 2014), resulting in 409,307 sites from 485,512 sites. The methylation level of a probe is represented by the “beta” value that is defined as a ratio of the intensity of methylation to the total intensity of probe.

In the 450K bead array, a probe measures DNA methylation level with either ‘type I’ assay or ‘type II’ assay. While the probe with type I assay employs two different bead types each for methylation and unmethylation with the same color channel, the probe with type II assay uses two different colors with one bead type to measure methylation and unmethylation status. Since the two types of probes measure DNA methylation level with different chemistries, it is necessary to adjust technical variation between them. We adopted a beta-mixture quantile dilation (BMIQ) method proposed by Teschendorff et al. (Teschendorff et al. 2013), to perform normalization of beta values within a sample.

### Computational analyses

Differentially methylated sites were identified between LTS- and STS-GBM groups by using R package, RUV-inverse for DNA methylation (Maksimovic et al. 2015). Briefly, this package evaluates beta values for a probe with generalized least squares regression after controlling batch effects based on negative control probes in the array. We required the following two conditions for significant differential methylation: i) p-value corrected for multiple testing by false discovery rate (FDR) is less than 0.01, ii) a site should be found in CpG context, excluding non-CpG methylation.

Pearson correlation coefficient was calculated between DNA methylation level (beta) for a given site and RNA expression level (log2 transformation of FPKM+1) of the corresponding refSeq mRNA using 32 TCGA GBM samples that are available for both of RNA-seq and 450k bead array. When multiple sites were matched to a single gene, the maximum value of correlation coefficient was assigned for a gene. Pearson correlation coefficient was calculated only when standard deviation of beta values across 32 GBM samples is greater than 0.1 to focus the sites with enough variation of DNA methylation.

The list of somatic mutation in GBM was downloaded from TCGA for 135 samples that have both of whole exome sequencing and 450k bead array. In each sample, a somatic mutation is associated with a probe site if the mutation is found between 5kbp upstream and downstream a probe site.

Gene ontology (GO) analysis for a given set of sites on 450k bead array was performed adjusting for the selection bias of genes inherent from non-uniform distribution of probes across genome. We took a weighted resampling approach similar with the previous report (Young et al. 2010). Specifically, 1000 times of random selection of as many genes as the identified genes was performed where a gene has selection weights according to the number of probes assigned to a gene. Statistical significance of a GO term was determined by comparing the number of genes associated with the GO term between the given gene set and the randomly-selected gene sets.

### Open source datasets and annotation

The probes on the methylation array were annotated with R package, “IlluminaHumanMethylation450kanno.ilmn12.hg19”, except for assignment of genes that were determined by the software ANNOVAR (Wang et al. 2010) using NCBI reference genes (Refseq). Note that the probes were grouped into four categories according to the distance from the closest CGI in the UCSC annotation: i) “island” if a probe is inside CGI, ii) “shore” if a probe is within 2000 bases, iii) “shore” if a probe is between 2000 and 4000 bases, iv) “open sea” otherwise. Further categories were made by distinguishing 5’ and 3’ direction from the closest CGI denoted by ‘N_’ and ‘S_’ respectively such as ‘N_shore’ and ‘S_shore’. The probes were also grouped according to the distance from transcription start sites (TSS) of the NCBI reference genes: i) “TSS1500” if a probe is within 200-1500 bases upstream TSS, ii) “TSS200” if a probe is between 0 and 200 bases upstream TSS, iii) “1st exon” if a probe is associated with first exon, iv) “Far” otherwise. If a probe has multiple categories, the following priority is applied to determine its single category: 1st Exon>TSS200>TSS1500>Far.

We obtained epigenetic annotation of human genomes from the ENCODE project (Consortium 2012; Sloan et al. 2016), including the DNaseI Hypersensitivity clusters in 125 cell types and the transcription factor binding sites of 161 factors for 91 cell types and the regions of ChIP
enrichment for 12 histone modifications (H2az, H3K27ac, H3K27me3, H3K36me3, H3K4me1, H3K4me2, H3K4me3, H3K79me2, H3K9ac, H3K9me1, H3K9me3, H4K20me1). The results from uniform processing by ENCODE Analysis Working Group were downloaded and manually parsed for further analyses. Also, we obtained H3K9me3 ChIP-seq and 450k bead array datasets of H1 from the ENCODE project.

The Cancer Genome Atlas (TCGA) datasets were downloaded for the level 3 data of RNA sequencing (RNA-seq), Illumina Infinium HumanMethylation450 BeadChip and exome sequencing of available GBM samples: FPKM, beta and the compiled list of somatic mutation sites respectively.

For U87MG, a GBM cell line, DNA methylation profiled by 450K bead array was obtained from ENCODE project. ChIP-seq peak (q-value<0.05) of H3K27ac was provided from Lin et al. 2012 (GSE36354).

## Acknowledgements

We appreciate for ENCODE Consortium and the ENCODE production laboratories generating the GBM dataset. Also, we should note that the results here are in part based upon data generated by the TCGA Research Network: http://cancergenome.nih.gov/.” This research was supported by the Basic Science Research Program through the National Research Foundation of Korea (NRF), funded by the Ministry of Education (NRF-2015-R1D1A1A09057171) in Korea, and the Seoul National University Hospital Research Fund (3020180010). We like to thank Min Jung Park, R.N. for the support in summarization of clinical data and sample management.

---

**Supplementary table 1.**
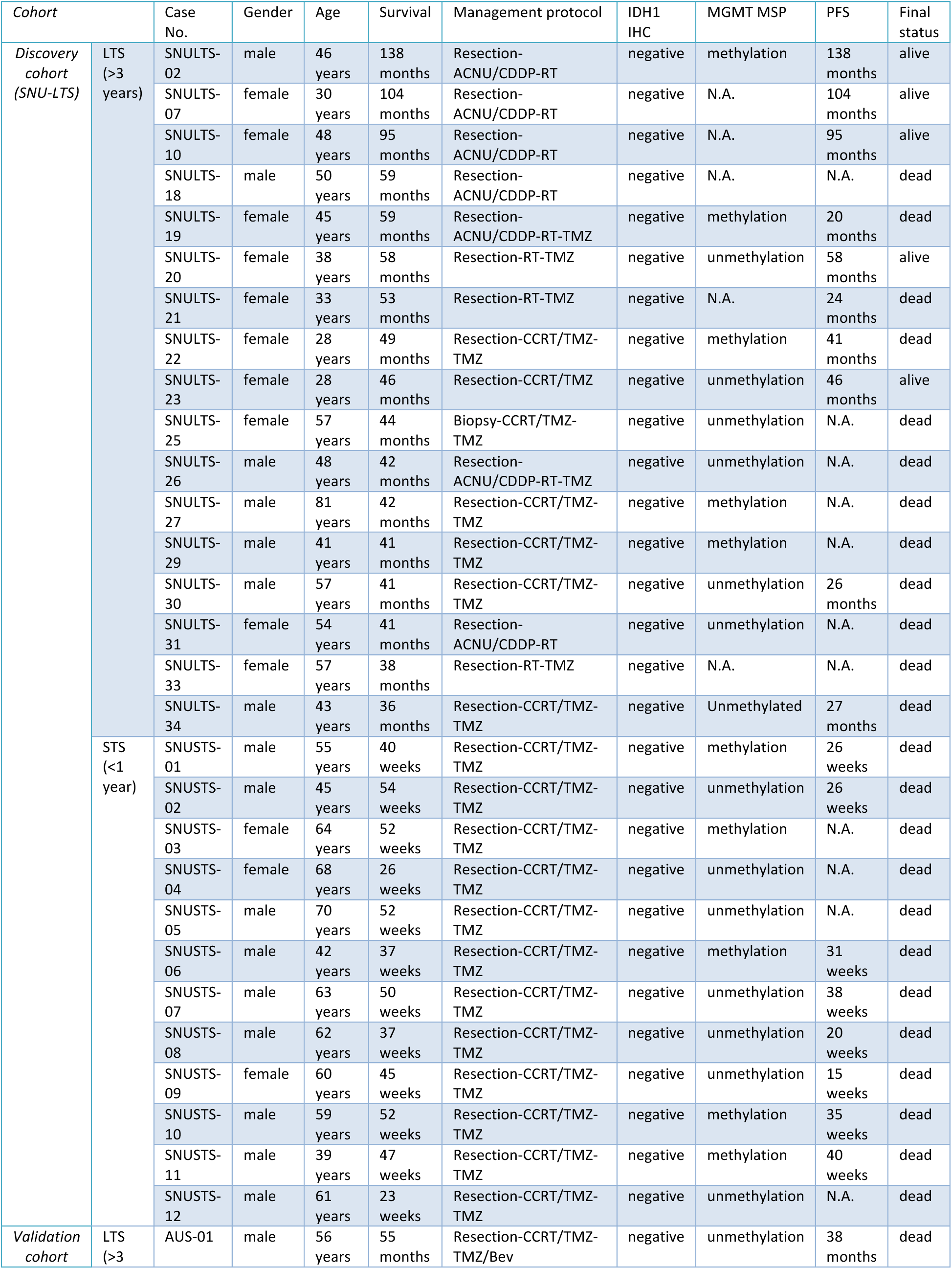
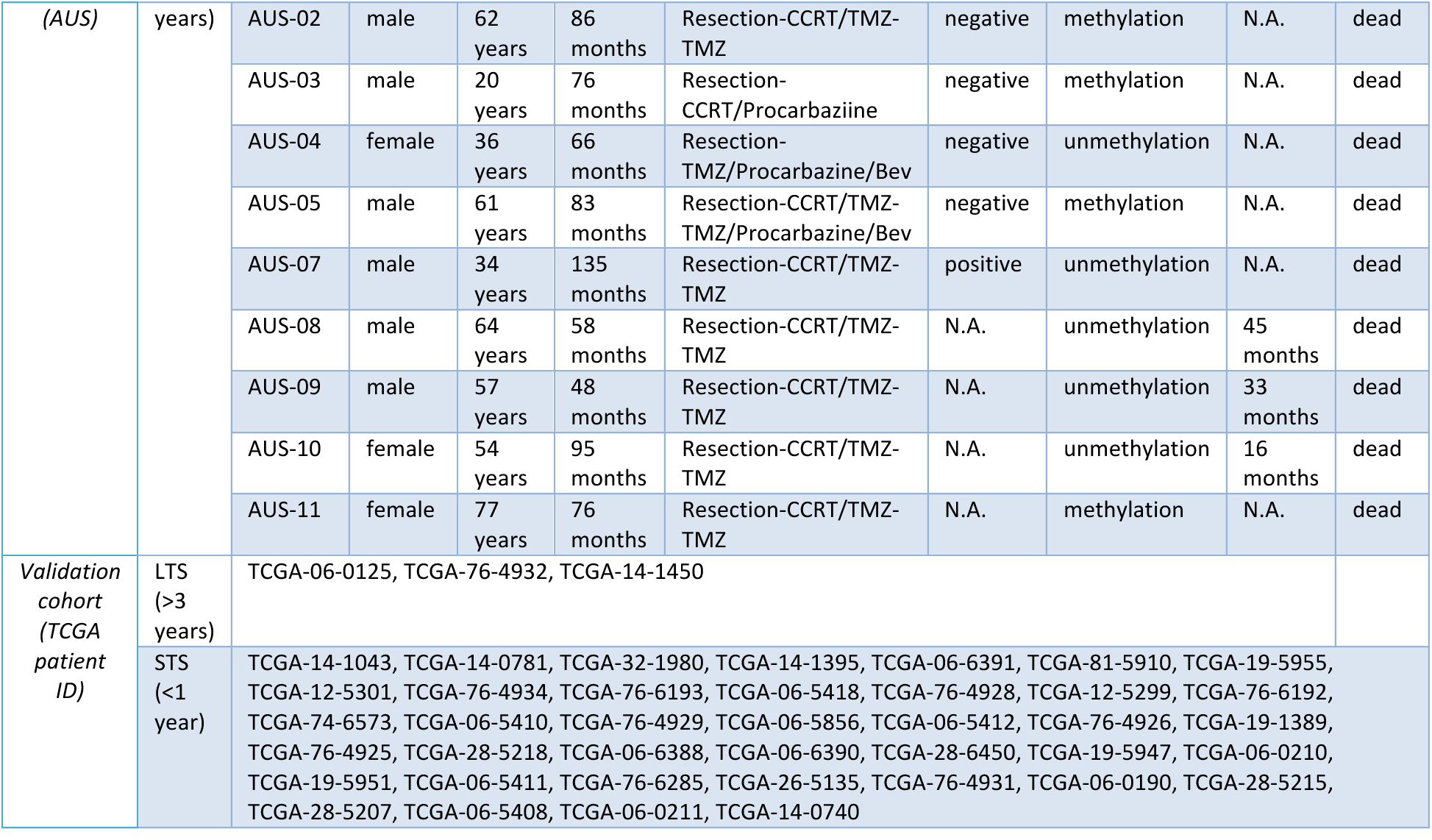
Sample information. Cohort: class of patient sample group (LTS; long-term survivor, STS; short-term survivor), Case No.: case-specific ID, Gender: male or female, Age: age at diagnosis in years, Diagnosis: histological diagnosis, Survival: survival period between first operation and death, IDH1 IHC: immunohistochemistry result of IDH1, MGMT MSP: MGMT methylation specific PCR, PFS: progression free survival, Final status: survival status at the last follow-up. For TCGA cohort, TCGA patient IDs are provided for reference.

**Supplementary table 2.** Gene Ontology (GO) results of the hyper-methylated sites. No: number, ID: GO ID, Category: GO category (BP: Biological Process, CC: Cellular Component, MF: Molecular Function), Description: GO Term description, N: the number of genes that are mapped in the 450K bead array in the GO term, Intersection: the number of genes that have associations with the selected hypermethylated sites among the genes in the previous column “N”, P-value: P-value, FDR: False Discovery Rate.

| No | ID | Category | Description | N | Intersection | P-value | FDR |
| --- | --- | --- | --- | --- | --- | --- | --- |
| 1 | GO:0015630 | CC | The part of the cytoskeleton (the internal framework of a cell) composed of microtubules and associated proteins. | 111 | 22 | 0 | 0.000 |
| 2 | GO:0008283 | BP | The multiplication or reproduction of cells, resulting in the expansion of a cell population. | 272 | 51 | 0.001 | 0.009 |
| 3 | GO:0006139 | BP | Any cellular metabolic process involving nucleobases, nucleosides, nucleotides and nucleic acids. | 24 | 8 | 0.002 | 0.018 |
| 4 | GO:0050680 | BP | Any process that stops, prevents or reduces the rate or extent of epithelial cell proliferation. | 43 | 13 | 0.002 | 0.018 |
| 5 | GO:0071364 | BP | Any process that results in a change in state or activity of a cell (in terms of movement, secretion, enzyme production, gene expression, etc.) as a result of an epidermal growth factor stimulus. | 21 | 8 | 0.002 | 0.018 |
| 6 | GO:0015629 | CC | The part of the cytoskeleton (the internal framework of a cell) composed of actin and associated proteins. Includes actin cytoskeleton-associated complexes. | 174 | 34 | 0.002 | 0.018 |
| 7 | GO:0016338 | BP | The attachment of one cell to another cell via adhesion molecules that do not require the presence of calcium for the interaction. | 14 | 6 | 0.003 | 0.027 |
| 8 | GO:0048008 | BP | The series of molecular signals generated as a consequence of a platelet-derived growth factor receptor binding to one of its physiological ligands. | 25 | 8 | 0.003 | 0.027 |
| 9 | GO:0000149 | MF | Interacting selectively and non-covalently with a SNARE (soluble N-ethylmaleimide-sensitive factor attached protein receptor) protein. | 36 | 10 | 0.003 | 0.027 |
| 10 | GO:0030041 | BP | Assembly of actin filaments by the addition of actin monomers to a filament. | 16 | 6 | 0.004 | 0.035 |
| 11 | GO:0045662 | BP | Any process that stops, prevents, or reduces the frequency, rate or extent of myoblast differentiation. A myoblast is a mononucleate cell type that, by fusion with other myoblasts, gives rise to the myotubes that eventually develop into skeletal muscle fibers. | 14 | 7 | 0.004 | 0.035 |
| 12 | GO:0006995 | BP | Any process that results in a change in state or activity of a cell (in terms of movement, secretion, enzyme production, gene expression, etc.) as a result of deprivation of nitrogen. | 12 | 5 | 0.005 | 0.043 |
| 13 | GO:0004843 | MF | Catalysis of the thiol-dependent hydrolysis of a peptide bond formed by the C-terminal glycine of ubiquitin. | 59 | 12 | 0.005 | 0.043 |
| 14 | GO:0005515 | MF | Interacting selectively and non-covalently with any protein or protein complex (a complex of two or more proteins that may include other nonprotein molecules). | 6498 | 767 | 0.005 | 0.043 |
| 15 | GO:0006665 | BP | The chemical reactions and pathways involving sphingolipids, any of a class of lipids containing the long-chain amine diol sphingosine or a closely related base (a sphingoid). | 64 | 14 | 0.006 | 0.051 |
| 16 | GO:0016328 | CC | The portion of the plasma membrane at the lateral side of the cell. In epithelial cells, lateral plasma membranes are on the sides of cells which lie at the interface of adjacent cells. | 39 | 11 | 0.006 | 0.051 |
| 17 | GO:0042832 | BP | Reactions triggered in response to the presence of a protozoan that act to protect the cell or organism. | 10 | 4 | 0.007 | 0.059 |
| 18 | GO:0090314 | BP | Any process that increases the frequency, rate or extent of the process of directing proteins towards a membrane, usually using signals contained within the protein. | 19 | 6 | 0.008 | 0.066 |
| 19 | GO:0000979 | MF | Interacting selectively and non-covalently with the regulatory region composed of the transcription start site and binding sites for transcription factors of the RNA polymerase II basal transcription machinery. | 42 | 13 | 0.008 | 0.066 |
| 20 | GO:0003774 | MF | Catalysis of the generation of force resulting either in movement along a microfilament or microtubule, or in torque resulting in membrane scission, coupled to the hydrolysis of a nucleoside triphosphate. | 35 | 8 | 0.008 | 0.066 |
| 21 | GO:0043548 | MF | Interacting selectively and non-covalently with a phosphatidylinositol 3-kinase, any enzyme that catalyzes the addition of a phosphate group to an inositol lipid at the 3' position of the inositol ring. | 19 | 7 | 0.009 | 0.074 |
| 22 | GO:0031492 | MF | Interacting selectively and non-covalently with the DNA portion of a nucleosome. | 25 | 7 | 0.01 | 0.081 |
| 23 | GO:0016579 | BP | The removal of one or more ubiquitin groups from a protein. | 45 | 10 | 0.011 | 0.088 |
| 24 | GO:0043388 | BP | Any process that increases the frequency, rate or extent of DNA binding. DNA binding is any process in which a gene product interacts selectively with DNA (deoxyribonucleic acid). | 20 | 7 | 0.011 | 0.088 |
| 25 | GO:0044804 | BP | A selective form of autophagy, by which damaged or non-essential parts of the nucleus, or even an entire nucleus is degraded. | 15 | 5 | 0.012 | 0.095 |
| 26 | GO:0019898 | CC | The component of a membrane consisting of gene products and protein complexes that are loosely bound to one of its surfaces, but not integrated into the hydrophobic region. | 70 | 14 | 0.012 | 0.095 |

**Supplementary table 3.** Gene Ontology (GO) results of the hypo-methylated sites. No: number, ID: GO ID, Category: GO category (BP: Biological Process, CC: Cellular Component, MF: Molecular Function), Description: GO Term description, N: the number of genes that are mapped in the 450K bead array in the GO term, Intersection: the number of genes that have associations with the selected hypo-methylated sites among the genes in the previous column “N”, P-value: P-value, FDR: False Discovery Rate.

| No | ID | Category | Description | N | Intersection | P-value | FDR |
| --- | --- | --- | --- | --- | --- | --- | --- |
| 1 | GO:0007186 | BP | A series of molecular signals that proceeds with an activated receptor promoting the exchange of GDP for GTP on the alpha-subunit of an associated heterotrimeric G-protein complex. The GTP-bound activated alpha-G-protein then dissociates from the beta- and gamma-subunits to further transmit the signal within the cell. The pathway begins with receptor-ligand interaction, or for basal GPCR signaling the pathway begins with the receptor activating its G protein in the absence of an agonist, and ends with regulation of a downstream cellular process, e.g. transcription. The pathway can start from the plasma membrane, Golgi or nuclear membrane (PMID:24568158 and PMID:16902576). | 609 | 297 | 0 | 0.000 |
| 2 | GO:0007608 | BP | The series of events required for an organism to receive an olfactory stimulus, convert it to a molecular signal, and recognize and characterize the signal. Olfaction involves the detection of chemical composition of an organism's ambient medium by chemoreceptors. This is a neurological process. | 176 | 91 | 0 | 0.000 |
| 3 | GO:0043928 | BP | The chemical reactions and pathways resulting in the breakdown of the transcript body of a nuclear-transcribed mRNA that occurs when the ends are not protected by the 3'-poly(A) tail. | 17 | 13 | 0 | 0.000 |
| 4 | GO:0050911 | BP | The series of events involved in the perception of smell in which an olfactory chemical stimulus is received and converted into a molecular signal. | 301 | 136 | 0 | 0.000 |
| 5 | GO:0004930 | MF | Combining with an extracellular signal and transmitting the signal across the membrane by activating an associated G-protein; promotes the exchange of GDP for GTP on the alpha subunit of a heterotrimeric G-protein complex. | 436 | 213 | 0 | 0.000 |
| 6 | GO:0004984 | MF | Combining with an odorant and transmitting the signal from one side of the membrane to the other to initiate a change in cell activity in response to detection of smell. | 301 | 136 | 0 | 0.000 |
| 7 | GO:0005549 | MF | Interacting selectively and non-covalently with an odorant, any substance capable of stimulating the sense of smell. | 77 | 37 | 0 | 0.000 |
| 8 | GO:0050482 | BP | The controlled release of arachidonic acid from a cell or a tissue. | 11 | 11 | 0.003 | 0.073 |
| 9 | GO:0015459 | MF | Modulates the activity of a potassium channel. | 26 | 24 | 0.003 | 0.073 |
| 10 | GO:0006103 | BP | The chemical reactions and pathways involving oxoglutarate, the dianion of 2-oxoglutaric acid. It is a key constituent of the TCA cycle and a key intermediate in amino-acid metabolism. | 13 | 10 | 0.004 | 0.086 |
| 11 | GO:0030101 | BP | The change in morphology and behavior of a natural killer cell in response to a cytokine, chemokine, cellular ligand, or soluble factor. | 11 | 10 | 0.004 | 0.086 |
| 12 | GO:0046872 | MF | Interacting selectively and non-covalently with any metal ion. | 1217 | 670 | 0.004 | 0.086 |
| 13 | GO:0030032 | BP | Formation of a lamellipodium, a thin sheetlike extension of the surface of a migrating cell. | 28 | 25 | 0.005 | 0.094 |
| 14 | GO:0003677 | MF | Any molecular function by which a gene product interacts selectively and non-covalently with DNA (deoxyribonucleic acid). | 834 | 433 | 0.005 | 0.094 |
| 15 | GO:0000288 | BP | A major pathway of degradation of nuclear-transcribed mRNAs that proceeds through a series of ordered steps that includes poly(A) tail shortening and that can regulate mRNA stability. | 34 | 21 | 0.006 | 0.099 |

## References

Amelot A, De Cremoux P, Quillien V, Polivka M, Adle-Biassette H, Lehmann-Che J, Francoise L, Carpentier AF, George B, Mandonnet E et al. 2015. IDH-Mutation Is a Weak Predictor of Long-Term Survival in Glioblastoma Patients. PloS one 10: e0130596.

Brennan CW, Verhaak RG, McKenna A, Campos B, Noushmehr H, Salama SR, Zheng S, Chakravarty D, Sanborn JZ, Berman SH et al. 2013. The somatic genomic landscape of glioblastoma. Cell 155: 462-477.

Burton EC, Lamborn KR, Feuerstein BG, Prados M, Scott J, Forsyth P, Passe S, Jenkins RB, Aldape KD. 2002a. Genetic aberrations defined by comparative genomic hybridization distinguish long-term from typical survivors of glioblastoma. Cancer Res 62: 6205-6210.

Burton EC, Lamborn KR, Forsyth P, Scott J, O’Campo J, Uyehara-Lock J, Prados M, Berger M, Passe S, Uhm J et al. 2002b. Aberrant p53, mdm2, and proliferation differ in glioblastomas from long-term compared with typical survivors. Clin Cancer Res 8: 180-187.

Charlet J, Duymich CE, Lay FD, Mundbjerg K, Dalsgaard Sorensen K, Liang G, Jones PA. 2016. Bivalent Regions of Cytosine Methylation and H3K27 Acetylation Suggest an Active Role for DNA Methylation at Enhancers. Molecular cell 62: 422-431.

Consortium TEP. 2012. An integrated encyclopedia of DNA elements in the human genome. Nature 489: 57.

Ehrlich M. 2009. DNA hypomethylation in cancer cells. Epigenomics 1: 239-259.

Geisenberger C, Mock A, Warta R, Rapp C, Schwager C, Korshunov A, Nied AK, Capper D, Brors B, Jungk C et al. 2015. Molecular profiling of long-term survivors identifies a subgroup of glioblastoma characterized by chromosome 19/20 co-gain. Acta neuropathologica 130: 419-434.

Gerber NK, Goenka A, Turcan S, Reyngold M, Makarov V, Kannan K, Beal K, Omuro A, Yamada Y, Gutin P et al. 2014. Transcriptional diversity of long-term glioblastoma survivors. Neurooncology 16: 1186-1195.

Hartmann C, Hentschel B, Simon M, Westphal M, Schackert G, Tonn JC, Loeffler M, Reifenberger G, Pietsch T, von Deimling A et al. 2013. Long-term survival in primary glioblastoma with versus without isocitrate dehydrogenase mutations. Clin Cancer Res 19: 5146-5157.

Jones PA. 2012. Functions of DNA methylation: islands, start sites, gene bodies and beyond. Nature reviews Genetics 13: 484-492.

Krex D, Klink B, Hartmann C, von Deimling A, Pietsch T, Simon M, Sabel M, Steinbach JP, Heese O, Reifenberger G et al. 2007. Long-term survival with glioblastoma multiforme. Brain 130: 2596-2606.

Lin CY, Lovén J, Rahl PB, Paranal RM, Burge CB, Bradner JE, Lee TI, Young RA. 2012. Transcriptional Amplification in Tumor Cells with Elevated c-Myc. Cell 151: 56-67.

Lu J, Cowperthwaite MC, Burnett MG, Shpak M. 2016. Molecular Predictors of Long-Term Survival in Glioblastoma Multiforme Patients. PloS one 11: e0154313.

Ma J, Hou X, Li M, Ren H, Fang S, Wang X, He C. 2015. Genome-wide methylation profiling reveals new biomarkers for prognosis prediction of glioblastoma. Journal of cancer research and therapeutics 11 Suppl 2: C212-215.

Mack SC, Hubert CG, Miller TE, Taylor MD, Rich JN. 2016. An epigenetic gateway to brain tumor cell identity. Nature neuroscience 19: 10-19.

Makova KD, Hardison RC. 2015. The effects of chromatin organization on variation in mutation rates in the genome. Nature reviews Genetics 16: 213-223.

Maksimovic J, Gagnon-Bartsch JA, Speed TP, Oshlack A. 2015. Removing unwanted variation in a differential methylation analysis of Illumina HumanMethylation450 array data. Nucleic Acids Res 43: e106.

Malzkorn B, Wolter M, Riemenschneider MJ, Reifenberger G. 2011. Unraveling the glioma epigenome: from molecular mechanisms to novel biomarkers and therapeutic targets. Brain Pathol 21: 619-632.

Martinez R, Schackert G, Yaya-Tur R, Rojas-Marcos I, Herman JG, Esteller M. 2007. Frequent hypermethylation of the DNA repair gene MGMT in long-term survivors of glioblastoma multiforme. Journal of neuro-oncology 83: 91-93.

Millward CP, Brodbelt AR, Haylock B, Zakaria R, Baborie A, Crooks D, Husband D, Shenoy A, Wong H, Jenkinson MD. 2016. The impact of MGMT methylation and IDH-1 mutation on long-term outcome for glioblastoma treated with chemoradiotherapy. Acta neurochirurgica 158: 1943-1953.

Mock A, Geisenberger C, Orlik C, Warta R, Schwager C, Jungk C, Dutruel C, Geiselhart L, Weichenhan D, Zucknick M et al. 2016. LOC283731 promoter hypermethylation prognosticates survival after radiochemotherapy in IDH1 wild-type glioblastoma patients. International journal of cancer 139: 424-432.

Morris TJ, Butcher LM, Feber A, Teschendorff AE, Chakravarthy AR, Wojdacz TK, Beck S. 2014. ChAMP: 450k Chip Analysis Methylation Pipeline. Bioinformatics 30: 428-430.

Nakagawa Y, Sasaki H, Ohara K, Ezaki T, Toda M, Ohira T, Kawase T, Yoshida K. 2017. Clinical and molecular prognostic factors for long-term survival of the patients with glioblastomas in a single institutional consecutive cohort. World neurosurgery doi:10.1016/j.wneu.2017.06.126.

Ostrom QT, Gittleman H, Xu J, Kromer C, Wolinsky Y, Kruchko C, Barnholtz-Sloan JS. 2016. CBTRUS Statistical Report: Primary Brain and Other Central Nervous System Tumors Diagnosed in the United States in 2009-2013. Neuro-oncology 18: v1-v75.

Parsons DW, Jones S, Zhang X, Lin JC-H, Leary RJ, Angenendt P, Mankoo P, Carter H, Siu IM, Gallia GL et al. 2008. An Integrated Genomic Analysis of Human Glioblastoma Multiforme. Science 321: 1807.

Peng S, Dhruv H, Armstrong B, Salhia B, Legendre C, Kiefer J, Parks J, Virk S, Sloan AE, Ostrom QT et al. 2017. Integrated genomic analysis of survival outliers in glioblastoma. Neurooncology 19: 833-844.

Prasanna P, Patel J, Partovi S, Madabhushi A, Tiwari P. 2016. Radiomic features from the peritumoral brain parenchyma on treatment-naive multi-parametric MR imaging predict long versus short-term survival in glioblastoma multiforme: Preliminary findings. European radiology doi:10.1007/s00330-016-4637-3.

Reifenberger G, Weber RG, Riehmer V, Kaulich K, Willscher E, Wirth H, Gietzelt J, Hentschel B, Westphal M, Simon M et al. 2014. Molecular characterization of long-term survivors of glioblastoma using genome- and transcriptome-wide profiling. International journal of cancer 135: 1822-1831.

Rideout WM, 3rd, Coetzee GA, Olumi AF, Jones PA. 1990. 5-Methylcytosine as an endogenous mutagen in the human LDL receptor and p53 genes. Science 249: 1288-1290.

Rose NR, Klose RJ. 2014. Understanding the relationship between DNA methylation and histone lysine methylation. Biochimica et biophysica acta 1839: 1362-1372.

Sarmiento JM, Mukherjee D, Black KL, Fan X, Hu JL, Nuno MA, Patil CG. 2016. Do Long-Term Survivor Primary Glioblastoma Patients Harbor IDH1 Mutations? Journal of neurological surgery Part A, Central European neurosurgery 77: 195-200.

Schuster-Bockler B, Lehner B. 2012. Chromatin organization is a major influence on regional mutation rates in human cancer cells. Nature 488: 504-507.

Scott JN, Rewcastle NB, Brasher PM, Fulton D, MacKinnon JA, Hamilton M, Cairncross JG, Forsyth P. 1999. Which glioblastoma multiforme patient will become a long-term survivor? A population-based study. Ann Neurol 46: 183-188.

Shinawi T, Hill VK, Krex D, Schackert G, Gentle D, Morris MR, Wei W, Cruickshank G, Maher ER, Latif F. 2013. DNA methylation profiles of long- and short-term glioblastoma survivors. Epigenetics 8: 149-156.

Shinojima N, Kochi M, Hamada J, Nakamura H, Yano S, Makino K, Tsuiki H, Tada K, Kuratsu J, Ishimaru Y et al. 2004. The influence of sex and the presence of giant cells on postoperative long-term survival in adult patients with supratentorial glioblastoma multiforme. J Neurosurg 101: 219-226.

Sloan CA, Chan ET, Davidson JM, Malladi VS, Strattan JS, Hitz BC, Gabdank I, Narayanan AK, Ho M, Lee BT et al. 2016. ENCODE data at the ENCODE portal. Nucleic Acids Research 44: D726-D732.

Smrdel U, Popovic M, Zwitter M, Bostjancic E, Zupan A, Kovac V, Glavac D, Bokal D, Jerebic J. 2016. Long-term survival in glioblastoma: methyl guanine methyl transferase (MGMT) promoter methylation as independent favourable prognostic factor. Radiology and oncology 50: 394-401.

Teschendorff AE, Marabita F, Lechner M, Bartlett T, Tegner J, Gomez-Cabrero D, Beck S. 2013. A beta-mixture quantile normalization method for correcting probe design bias in Illumina Infinium 450 k DNA methylation data. Bioinformatics 29: 189-196.

Turcan S, Rohle D, Goenka A, Walsh LA, Fang F, Yilmaz E, Campos C, Fabius AWM, Lu C, Ward PS et al. 2012. IDH1 mutation is sufficient to establish the glioma hypermethylator phenotype. Nature 483: 479.

Wang K, Li M, Hakonarson H. 2010. ANNOVAR: functional annotation of genetic variants from high-throughput sequencing data. Nucleic Acids Research 38: e164-e164.

Young MD, Wakefield MJ, Smyth GK, Oshlack A. 2010. Gene ontology analysis for RNA-seq: accounting for selection bias. Genome Biology 11: R14.

Zhang W, Yan W, You G, Bao Z, Wang Y, Liu Y, You Y, Jiang T. 2013. Genome-wide DNA methylation profiling identifies ALDH1A3 promoter methylation as a prognostic predictor in G-CIMP-primary glioblastoma. Cancer letters 328: 120-125.

